# *Clostridium difficile* alters the structure and metabolism of distinct cecal microbiomes during initial infection to promote sustained colonization

**DOI:** 10.1101/211516

**Authors:** Matthew L. Jenior, Jhansi L. Leslie, Vincent B. Young, Patrick D. Schloss

## Abstract

Susceptibility to *Clostridium difficile* infection is primarily associated with previous exposure to antibiotics, which compromise the structure and function of the gut bacterial community. Specific antibiotic classes correlate more strongly with recurrent or persistent *C. difficile* infection. As such, we utilized a mouse model of infection to explore the effect of distinct antibiotic classes on the impact that infection has on community-level transcription and metabolic signatures shortly following pathogen colonization and how those changes may associate with persistence of *C. difficile*. Untargeted metabolomic analysis revealed that *C. difficile* infection had significantly larger impacts on the metabolic environment across cefoperazone and streptomycin-pretreated mice, which become persistently colonized compared to clindamycin-pretreated mice where infection quickly became undetectable. Through metagenome-enabled metatranscriptomics we observed that transcripts for genes associated with carbon and energy acquisition were greatly reduced in infected animals, suggesting those niches were instead occupied by *C. difficile*. Furthermore, the largest changes in transcription were seen in the least abundant species indicating that *C. difficile* may “attack the loser” in gut environments where sustained infection occurs more readily. Overall, our results suggest that *C. difficile* is able to restructure the nutrient-niche landscape in the gut to promote persistent infection.

## Importance

*Clostridium difficile* has become the most common single cause of hospital-acquired infection over the last decade in the United States. Colonization resistance to the nosocomial pathogen is primarily provided by the gut microbiota, which is also involved in clearing the infection as the community recovers from perturbation. As distinct antibiotics are associated with different risk levels for CDI, we utilized a mouse model of infection with 3 separate antibiotic pretreatment regimes to generate alternative gut microbiomes that each allowed for *C. difficile* colonization but varied in clearance rate. To assess community-level dynamics, we implemented an integrative multi-omic approach that revealed that infection significantly changed many aspects of the gut community. The degree to which the community changed was inversely correlated with clearance during the first six days of infection, suggesting that *C. difficile* differentially modifies the gut environment to promote persistence. This is the first time metagenome-enabled metatranscriptomics have been employed to study the behavior of a host-associated microbiota in response to an infection. Our results allow for a previously unseen understanding of the ecology associated with *C. difficile* infection and provides groundwork for identification of context-specific probiotic therapies.

## Introduction

One of the many beneficial functions provided by the indigenous gut bacterial community is its ability to protect the host from infection by pathogens (1). This attribute, termed colonization resistance, is one of the main mechanisms that protect healthy individuals from the gastrointestinal pathogen *Clostridium difficile* (2–4). *C. difficile* infection (CDI) is responsible for most cases of antibiotic-associated colitis, a toxin-mediated diarrheal disease that has dramatically increased in prevalence over the last 10 years. There are an estimated 453,000 cases of CDI resulting in 29,000 deaths in the United States annually (5). Antibiotics are a major risk factor for CDI and are thought to increase susceptibility by disrupting the gut bacterial community structure; however, it is still unclear what specific changes to the microbiota contribute to this susceptibility (6, 7). Although most classes of antibiotics have been associated with initial susceptibility to CDI, fluoroquinolones, clindamycin, and cephalosporins are linked to increased risk of recurrent or persistent infection (8–10). This raises questions about the groups of bacteria that are differentially impacted by certain therapies and how these changes effect duration or severity of the infection.

Associations between the membership and functional capacity of the microbiota as measured by the metabolic output suggest that antibiotics increase susceptibility by altering the nutrient milieu in the gut to one that favors *C. difficile* metabolism (11–13). One hypothesis is that *C. difficile* colonization resistance is driven by competition for growth substrates by an intact community of metabolic specialists. This has been supported by animal model experiments over the past several decades (14–16). This line of reasoning has been carried through to the downstream restoration of colonization resistance with the application of fecal microbiota transplant (FMT). Although an individual’s microbiota may not return to its precise original state following FMT, it is hypothesized that the functional capacity of the new microbiota is able to outcompete *C. difficile* for resources and clear the infection (13, 17).

Leveraging distinct antibiotic treatment regimens in a murine model of CDI (18), we and others have shown that *C. difficile* adapts its physiology to the distinct cecal microbiomes that resulted from exposure to antibiotics (18, 19). We went on to show that *C. difficile* appears to adapt portions of its metabolism to fit alternative nutrient niche landscapes. As the diet of the mice remained unchanged, changes in the cecal metabolome are likely driven by the intestinal microbiota. Although it has been established that *C. difficile* colonizes these communities effectively, it is unknown whether the differences in the metabolic activity of communities following antibiotic treatment are impacted by *C. difficile* colonization or if they correlate with prolonged infection. Historically, it has been difficult to ascribe specific metabolic contributions to individual taxa within the microbiota during perturbations, especially within the context of a host. To address this limited understanding, we employed an integrative untargeted metabolomic and metagenome-enabled metatransciptomic approach to investigate specific responses to infection of the gut microbiota in a murine model of CDI. This high-dimensional analysis allowed us to not only characterize the metabolic output of the community, but to also identify which subgroups of bacteria were differentially active during mock infection and CDI. Our results supported the hypothesis that CDI was indeed associated with altered community-level gene transcription and metabolomic profile of susceptible environments. This effect was significantly more pronounced in communities where *C. difficile* was able to maintain colonization. This work highlights the need for increased appreciation of the differential, combined effects of antibiotics and CDI on the gut microbiota to develop more successful targeted therapies that eliminate *C. difficile* colonization.

## Results

### Distinct antibiotic pretreatments are associated with alternative community structures that are equally susceptible to initial *C. difficile* colonization, but differ in patterns of clearance

We have previously shown that when conventionally-reared SPF mice were pretreated with one of three different antibiotics (streptomycin, cefoperazone, and clindamycin; Table S1), each pretreatment was associated with altered patterns of *C. difficile* virulence factor expression (19). These antibiotics were chosen for not only the ability to to reduce *C. difficile* colonization resistance in a mouse model (18), but also for distinct and significant impacts on the structure and diversity of the cecal microbiota (Fig. 1A) (19). In each antibiotic pretreatment, we observed equally high levels of *C. difficile* colonization on the day after infection, however, over the subsequent 9 days the amount of *C. difficile* in the feces of clindamycin-pretreated mice were the only mice to fall below the limit of detection, while mice receiving the other pretreatments remained highly colonized (*p* = 0.01; Fig. 1A). We hypothesized that this occurred in the clindamycin-pretreated mice because the perturbed intestinal community occupied niche space that overlapped with that of *C. difficile*.

**Figure 1.**
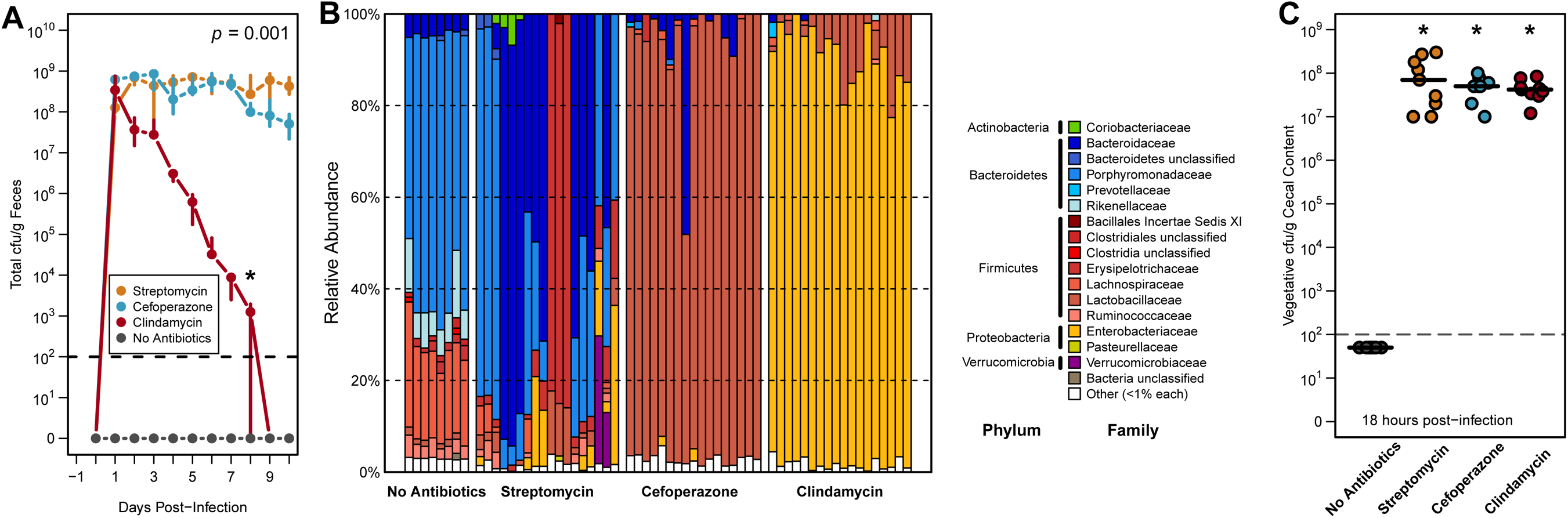
Distinct antibiotic pretreatments have differential impacts on *C. difficile* colonization and cecal microbiota community structure. **(A)** C. *difficile* 630 CFU in stool of infected mice following each antibiotic-pretreated group over 10 days of infection. Median and interquartile range are shown for each time point. Both cefoperzone and streptomycin pretreatments had more significantly detectable CFU on the final day of detectable CFU associated with clindamycin-pretreatment (*p* < 0.001). **(B)** Relative abundance of family-level OTU taxonomic classification in each pretreatment group from 16S rRNA gene sequencing. **(C)** Quantification of terminal vegetative *C. difficile* CFU in cecal content across 18 hour colonization models. Black lines indicate median values and each pretreatment group had significantly greater detectable CFU than no antibiotic controls. Significant differences in A were determined through permANOVA with dynamic time warping and in C were found by Wilcoxon rank-sum test with Benjamni-Hochberg correction when necessary. The limit of detection was used in place of undetectable values for statistical testing.

We chose to focus our remaining experiments on cecal samples collected 18 hours after infection to the assess behavior of *C. difficile* directly prior to the reduction in detectable *C. difficile*. This end point corresponded with a previous study where *C. difficile* reached maximum cecal vegetative cell load with few detectable spores (20). We also elected to examine cecal content because it was more likely to be a site of active bacterial metabolism compared to stool and would allow for an assessment of functional differences in the microbiota. At 18 hours after infection, we found that the communities remained highly differentiated from untreated controls as measured by 16S rRNA gene sequencing of the V4 region (Fig. 1B). The composition of streptomycin-pretreated communities was more variable between cages, but was generally enriched for members of the *Bacteroidetes* phylum. Cefoperazone and clindamycin-pretreated cecal communities were consistently dominated by members of the *Lactobacillaceae* and *Enterobacteriaceae* families, respectively. Despite variation in the community structures, there were no significant differences in the number of vegetative cells between any antibiotic- pretreatment group (Fig. 1C). All susceptible mice were colonized with ~1×10^8^ vegetative colony forming units (CFU) per gram of cecal content and untreated mice maintained *C. difficile* colonization resistance. We have also previously demonstrated that both *C. difficile* spore production and toxin activity differ between these pretreatment regimes (19). As both processes have been linked to environmental concentrations of specific growth nutrients (21), these results suggested that despite high initial *C. difficile* colonization the microbiomes across pretreatments may vary in available nutrients or profiles of competitors for those niches.

### Multiple biological signatures in the bacterial community and metabolome differentiated cecal microbiomes that remained colonized by *C. difficile* from those that did not

Pretreatment with antibiotics not only alters the structure of the resident microbiota, but also has a dramatic impact on the intestinal metabolome (11–13). To understand the ramifications each antibiotic had on the cecal metabolomic environment, we performed untargeted metabolomic analysis on the cecal contents that were also utilized in the 16S rRNA gene sequencing. We identified a total of 727 distinct metabolites. In combination with our 16S rRNA gene sequencing results, we first characterized the differences between the microbiomes (i.e. the microbiota, plus the associated metabolome) of the mock-infected animals to quantify possible drivers of communities that cleared the infection. To focus our analysis on ascertaining changes in discrete populations within the microbiota, we generated operational taxonomic units (OTUs) clustered at 97% similarity. We also removed all *C. difficile* 16S rRNA gene sequences, which represented an average of 2.113% sequencing reads across infection groups to eliminate its direct impact in downstream calculation. Using these methods we discovered that the Bray-Curtis dissimilarity of both the community structure (*p* < 0.001) and metabolome (*p* < 0.001) were significantly different between cleared and colonized groups during the early stages of infection (Fig. 2A & 2C). These results supported the hypothesis that the cecal environment created by clindamycin pretreatment was highly divergent from the other groups, and likely contributed to the clearance seen in the subsequent days.

**Figure 2.**
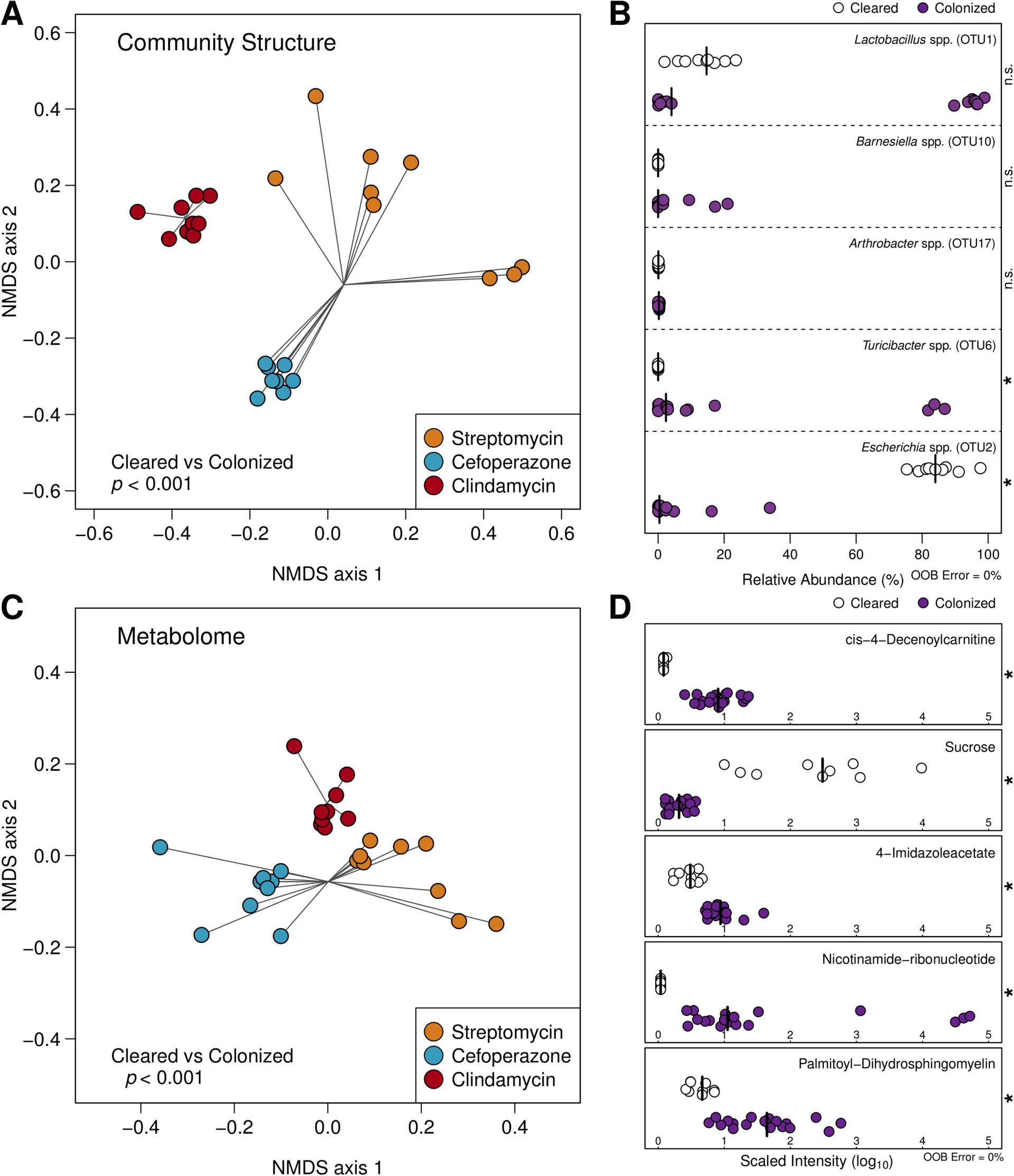
Significant differences in cecal community structure and metabolomes track with downstream *C. difficile* clearance across antibiotic pretreatment regimes. **(A)** NMDS ordination of Bray-Curtis distances of OTU relative abundances between mouse cecal communities that remained colonized by *C. difficile* and those that eventually cleared the infection. **(B)** Relative abundance of OTUs included the optimal model generated by AUCRF classifying the same groups as in panel A. Species-level identification was obtained using centroid representative sequences for each OTU. **(C)** NMDS ordination of Bray-Curtis distances using metabolite intensities between the aforementioned groups of animals. **(D)** Scaled intensity of metabolites included the optimal model generated by AUCRF classifying colonized and clearing mouse cecal microbiomes. Differences for ordinations in A & C were calculated using permANOVA. Optimal AUCRF models demonstrated 0% out of bag error, and significant differences in B & D were determined by Wilcoxon rank-sum test with Benjamni-Hochberg correction.

To identify the populations and metabolites that were associated with sustained colonization, we utilized Random Forest machine learning with cross validation to identify the smallest optimal subset of features that could successfully differentiate microbiomes that clear infection and and those that remain colonized (22). We identified a model with 5 OTUs that correctly classified all samples to their corresponding groups (Fig. 2B; Out-of-bag error=0%). Interestingly, these OTUs were not consistently abundant in antibiotic-pretreated communities. Similarly, when we used the same approach with the metabolomic data, we identified a model that used 5 metabolites that correctly differentiated the groups (Fig. 2D; Out-of-bag error=0%). Together these results further supported the hypothesis that the environment of the cecum, even early during infection, is distinct between groups that clear the infection and those that maintain *C. difficile* at high levels. Furthermore, results from machine learning analysis suggest that rare members of the communities had a disproportionate influence on the clearance patterns observed between pretreatment regimes and that changes in community structure may be less consistent than changes in the metatransciptome or metabolome.

### Amino-acid metabolism by *C. difficile* appears important for sustained colonization across susceptible environments

*C. difficile's* ability to metabolize amino acids via Stickland fermentation may be a critical nutrient niche that enables it to colonize some perturbed communities (23). We were curious whether this behavior was conserved across multiple distinct gut environments where *C. difficile* was able to colonize. We assessed the changes between the antibiotic-pretreated, mock-infected microbiomes and those of untreated, *C. difficile-resistant* animals. Not only were the relative abundances of Stickland fermentation substrates increased across susceptible environments, but several secondary bile acids, which have been shown to be negatively correlated with *C. difficile* susceptibility were significantly decreased (Fig. S1D; *p* < 0.001). Additionally, when we constructed a Random Forest classification model to differentiate the groups, we identified multiple members of the *Clostridia* which are capable of metabolizing amino acids for growth (24). The relative abundances of these populations were significantly lower in susceptible animals (Fig. S1B; *p* < 0.001). We also performed a similar analysis to investigate changes induced by *C. difficile* colonization itself in these susceptible conditions. Although CDI alone did not induce significant shifts in the global community structure or metabolome (Fig. S2 A & C; *p* = 0.185, 0.065), several features were able to discriminate infected and uninfected microbiomes with high accuracy. This analysis highlighted numerous growth substrates that are known for *C. difficile* in all pretreated mice including 6 Stickland substrates, 4 of which were proline conjugates, along with arabonate/xlyonate (Fig. S2D). Furthermore 5-aminovalerate, the most common end product of Stickland fermentation, was significantly increased during infection in almost all of the metabolomes. Inspection of these specific metabolites revealed that clindamycin pretreatment was only condition where both the inputs and outputs of Stickland fermentation were less abundant relative to the untreated mice (Fig. S3). These results strongly support Stickland fermentation as a primary nutritional strategy of *C. difficile* early in infection. Moreover, these data suggest that the degree to which the environment of the intestine is altered by infection may be linked to the ability of the pathogen to remain colonized.

### Infection corresponded with larger shifts in the metatranscriptomes of communities that allowed sustained *C. difficile* colonization

Despite the strong associations between bacterial community structure and the metabolome with colonization resistance, it was difficult to associate specific populations with changes in those metabolites that were associated with the duration of infection. To gain a more specific understanding of how the microbiota or *C. difficile* shaped the metabolic environment, we employed parallel metagenomic and metatranscriptomic shotgun sequencing of the samples collected from the cecal content of the mice used in the previous analyses. To achieve usable concentrations of bacterial mRNA after rRNA depletion, we had to pool the samples within each treatment and infection group. To establish confidence in the results of a pooled analysis, we calculated within-group sample variance among replicates using CFU, OTU relative abundance, and metabolomic relative abundance data (Table S3). These analyses revealed low levels of variance within control and experimental groups. Following sequencing, metagenomic reads from mock-infected cecal communities were assembled *de novo* into contigs and putative genes were identified resulting in 234,868 (streptomycin), 83,534 (cefoperazone), and 35,681 (clindamycin) open reading frames in each metagenome. Of these putative genes, 28.5% could be annotated to a known function based on the KEGG database, and many of these annotations were homologs to genes in species that were found in our dataset. Streptomycin pretreatment resulted in a significantly more diverse community than other groups based on 16S rRNA gene sequence data, so a more diverse metagenome was expected (Table S1). Supporting this prediction, 2408 unique functionally annotated genes were detected in the streptomycin pretreatment metagenome, at least 1163 more genes than were found in either the cefoperazone or clindamycin metagenomes (Fig. S4A-D). Metagenome-enabled mapping of the metatranscriptomic reads revealed that we were able to obtain informative depths of sequencing from across the metagenomic libraries (Fig. S4E-F). As expected, genes with any detectable transcript in any metatranscriptome were a subset of their corresponding metagenome. Metatranscriptomic read abundances were normalized to corresponding metagenomic coverage per gene to normalize for the abundance of the contributing bacterial taxa. This step was followed by a final subsampling of reads from each conditions to control for uneven sequencing effort and to identify genes with the largest changes in transcription relative to uninfected animals.

We hypothesized that the degree of change in the metatranscriptome corresponding with *C. difficile* colonization would reflect the shifts seen at in the metabolome. As disparate bacterial taxa possess vastly different metabolic capabilities and the antibiotic pretreatments induced distinct species profiles in each community, we tested our hypothesis by delineating the transcriptomic contributions of separate bacterial taxa within each metatranscriptome. Since many genes lack a specific functional annotation in KEGG but retain general taxonomic information, we continued the analysis at the genus level of classification for all genes contributed to each metagenome. Using this approach, we directly compared the normalized transcript abundances for each gene between infected and uninfected states for each antibiotic pretreatment and calculated the Spearman correlation to identify distinct patterns of transcription (Fig. 3). This resulted in 2473 genes that had an average distance of 2.545 units of deviation (UD) associated with streptomycin-pretreatment, 2930 genes at an average distance of 3.854 UD with cefoperazone-pretreatment, and only 727 genes at an average distance of 2.414 UD with clindamycin-pretreatment. Overall, the clindamycin pretreatment was associated with the fewest transcription outliers between uninfected and infection conditions compared with those of the other antibiotic groups. This suggested that the degree to which the metatranscriptome was altered by infection corresponded to prolonged colonization.

**Figure 3.**
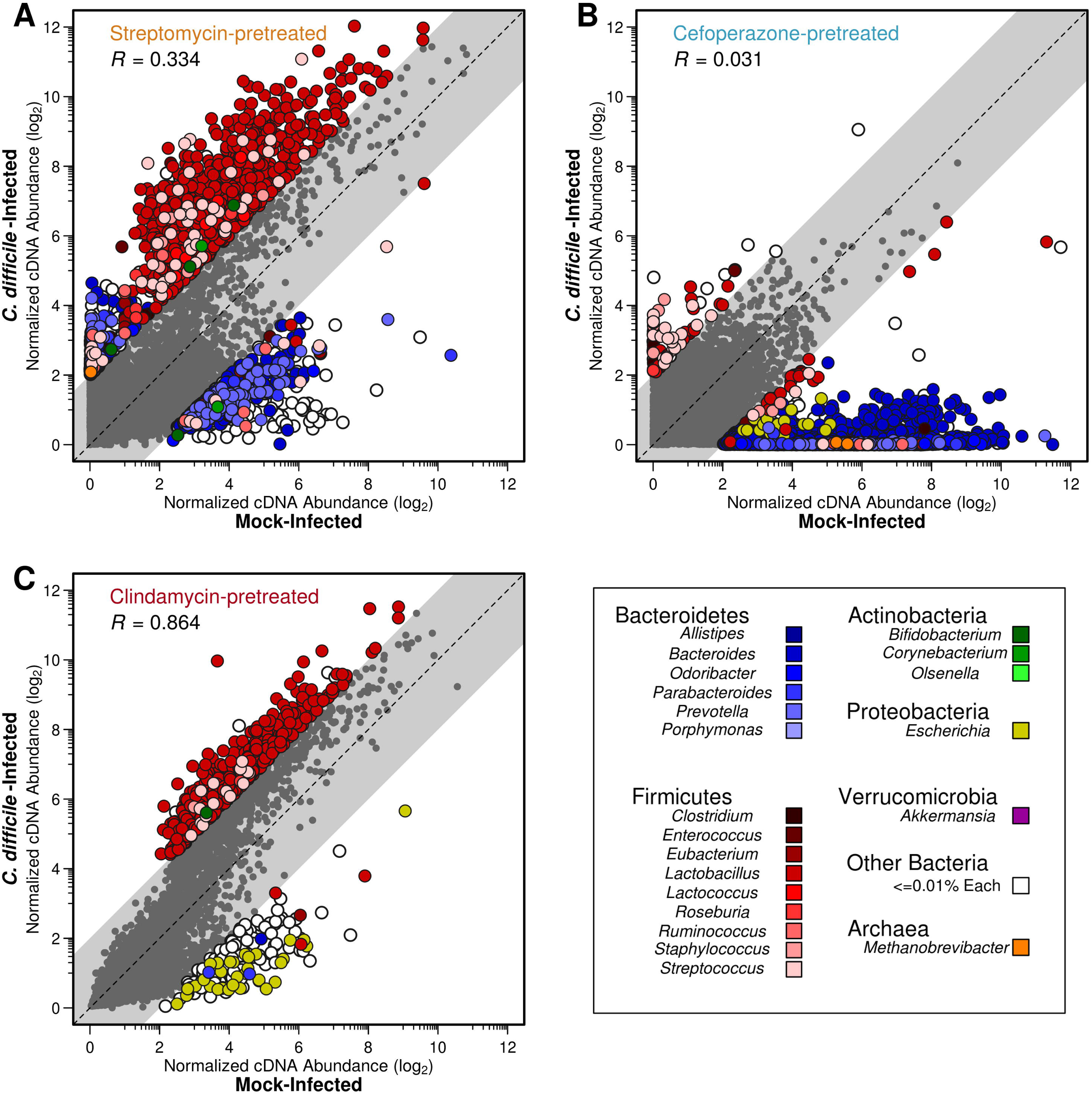
*C. difficile* colonization alters gene transcription of taxonomic groups differentially between antibiotic pretreatments. Each point represents a unique gene from the respective metagenomic assembly. Coordinates were determined by the log_2_-transformed difference in transcription level between *C. difficile-infected* and mock-infected conditions for each gene. Outliers were defined using linear correlation and a squared residual cutoff of 2. Distance of outliers to the x=y line were also calculated and represented in unites of deviation or UD. The coloring of each point indicates the genus that the transcript originated from and the and the gray points denote those genes with consistent transcription levels between conditions as defined by outlier analysis. Antibiotic pretreatments; **(A)** Streptomycin-pretreated, **(B)** Cefoperazone-pretreated, and **(C)** Clindamycin- pretreated.

This analysis also revealed that outlier genes originated in underrepresented genera. In streptomycin-pretreated mice, 937 genes belonging to *Lactobacillus* that higher transcription during *C. difficile* infection; *Lactobacillus* accounted for 0.42% of the 16S rRNA gene sequences (Fig. 3A). In cefoperazone-pretreated mice, 2290 genes belonging to *Bacteroides* had lower transcription during *C. difficile* infection; *Bacteroides* accounted for 1.49% of the 16S rRNA gene sequences (Fig. 3B). A consistent trend in streptomycin and cefoperazone-pretreated mice was an overrepresentation of highly transcribed genes from genera belonging to *Bacteroidetes* during mock infection. The metatransciptomes among mice from both of these pretreatment conditions poorly correlated between mock and infected conditions, indicating a high degree of change induced by *C. difficile* colonization (*R* = 0.334 & *R* = 0.031). In clindamycin-pretreated mice the largest difference in transcription was for 510 *Lactobacillus* genes with increased transcription during CDI; *Lactobacillus* accounted for 2.7% of the 16S rRNA gene sequences (Fig. 3C). Infected and uninfected metatranscriptomes from mice pretreated with clindamycin were more strongly correlated with each other than either of the other pretreatments (*R* = 0.864). This suggests that although *C. difficile* altered the streptomycin and cefoperazone- pretreated communities in which it was able to remain stably colonized, it had minimal impact on the clindamycin-pretreated community in which it was not able to remain colonized.

### Largest changes in metatranscriptomes in response to infection were concentrated in the minority taxa of each pretreatment group

To explore the observation that rare taxa were responsible for the largest differences in transcription in response to infection, we tabulated the absolute difference between mock and *C. difficile* infected transcriptomes for each genus in each antibiotic pretreatment. We further normalized these values for the number of genes detected in each genus to adjust for genera that were more successfully assembled or annotated and we eliminated genera where less than 50 genes were detected in the metatranscriptome. Taxa were then stratified into categories based on their relative abundance in each community from 16S rRNA gene sequencing (Fig. 4). This revealed that most change occurred among the rare genera and that the degree of change was inversely correlated with sustained colonization. To this point, minority metatranscriptomic absolute differences were significantly reduced in clindamycin pretreatment (*p* < 0.001). Additionally, the proportion of taxa in the lowest relative abundance bracket was similar across pretreatment groups (~88.9%). As a corollary, we predicted that the majority of unique genes or metabolic potential was held within this minority, and when following quantification this proved to be the case (Table S4). As a consequence, the downstream impacts on functionality may affect a disproportionately large effect on the overall environment of the intestine as a function of its collective metabolism.

**Figure 4.**
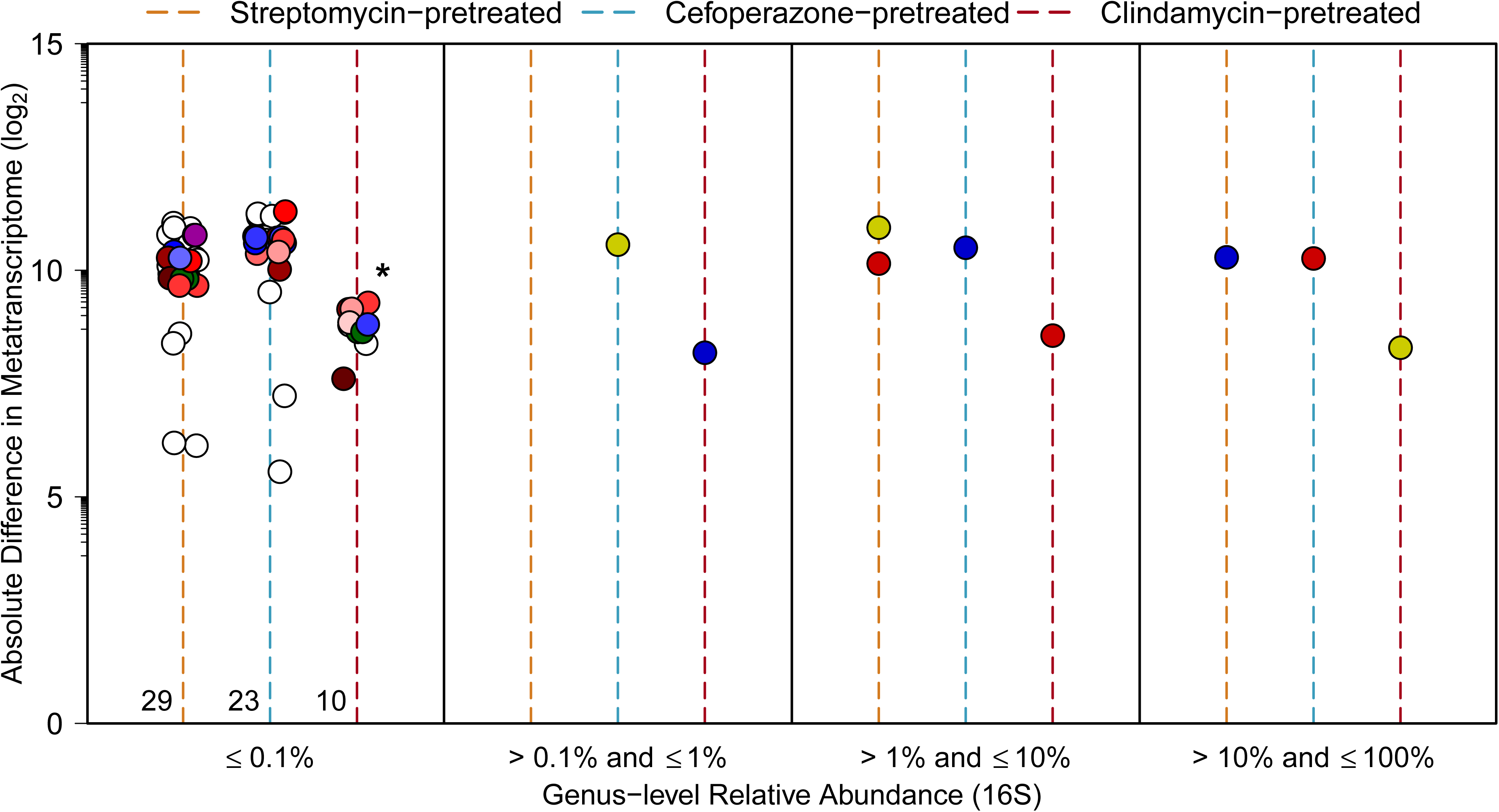
A majority of metatranscriptomic changes are focused within minority members of each microbiota. Absolute difference in metatranscriptomic reads contributed by each genus in pretreatments between mock and *C. difficile*-infected conditions. Colored lines denoted antibiotic pretreatment. Each point represents all transcript contributed by that genus in each pretreatment group. Numbers at the base of pretreatment lines in the first panel represent the quantity of genera in each group as some points are obscured.

### Altered transcription within minority taxa favors reduced nutrient competition with *C. difficile* in communities that permitted sustained colonization

Based on our metabolomic and metatranscriptomic results, we hypothesized that pathways with the greatest differences between mock and *C. difficile-infected* mice would be related to catabolism of metabolites that *C. difficile* could use for for growth. To assess these changes, we identified those annotated transcripts that were associated with genera that represented less than 0.1% of the community as measured with our 16S rRNA gene sequence data (Fig. 5). This resulted in the identification of 585 genes that were differentially transcribed between clindamycin-pretreated mice and the streptomycin and cefoperazone-pretreated mice. From this group of genes we filtered the collection to identify those genes that were unique to either the clindamycin-pretreated mice or the streptomycin and cefoperazone-pretreated mice. Finally, we limited our analysis to those genes that were meaningfully different between the mock and *C. difficile-infected* groups in each antibiotic pretreatment group. This resulted in 34 genes from 11 pathways. These genes and pathways were primarily involved in simple carbohydrate-containing molecule acquisition/utilization (Fig. 5). Interestingly, many of these genes had decreased transcription during infection compared to mock-infected controls. At the pathway-level, many genes associated with galactose and amino sugar acquisition (both *C. difficile* growth substrates) were reduced during infection in both streptomycin and cefoperazone- pretreated mice. Conversely, pathways uniquely associated with clindamycin-pretreated communities were related to the metabolism of a diverse array of carbon sources, which may indicate ineffective competition by *C. difficile* with this community for any particular growth substrate. Our results indeed suggest that *C. difficile* colonization induces a shift transcriptional activity for a minority subset of species, possibly in an effort to segregate a desired nutrient niche, prior to the introduction of the hallmark disease phenotypes associated with CDI.

**Figure 5.**
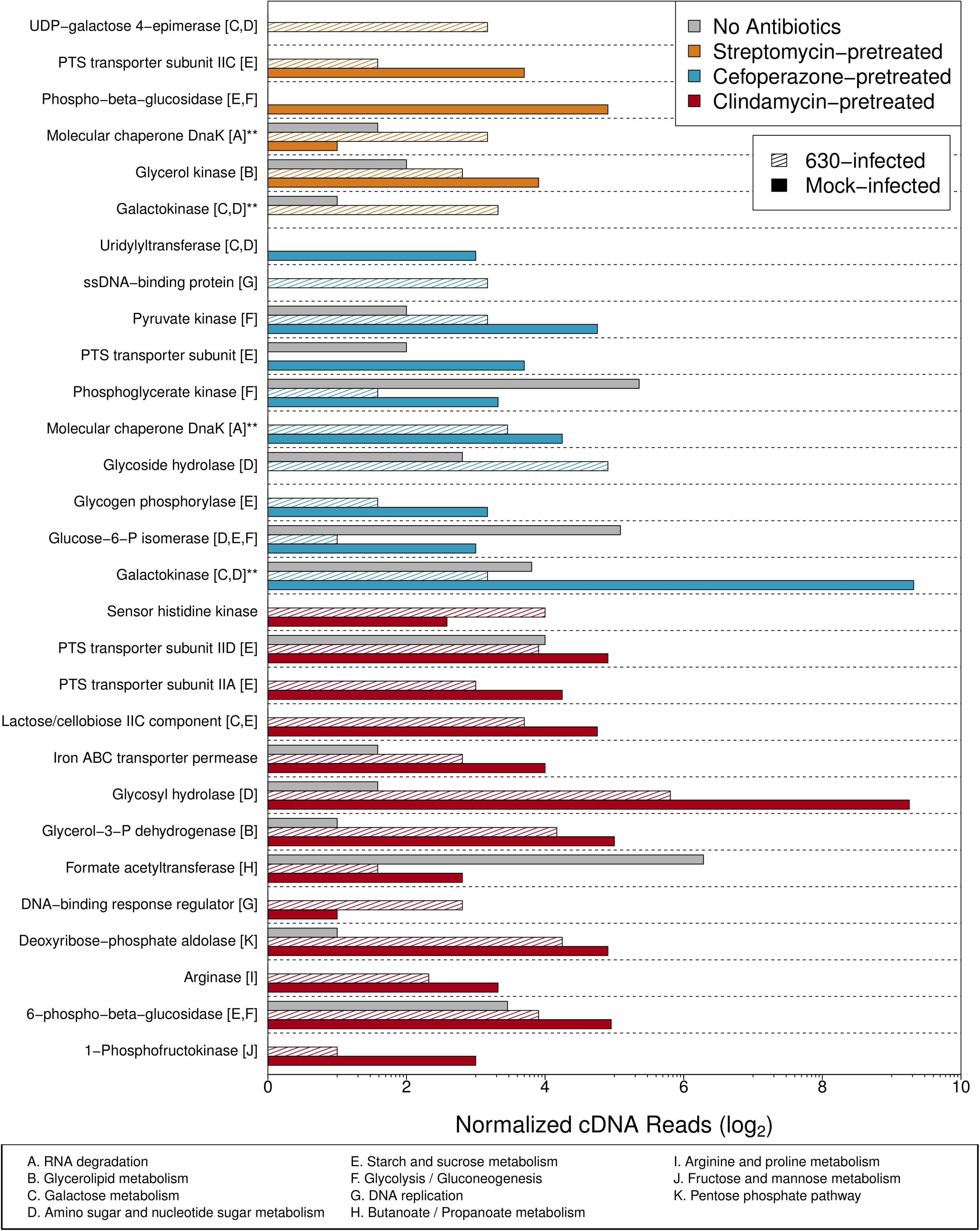
Metatranscriptomic changes due to infection in certain metabolic pathways are overrepresented in the minority taxa. Log_2_ metagenome-normalized cDNA abundances for genes with differential transcription during infection belonging to genera that had a relative abundance greater than 0.1%. Double asterisks denotes genes shared between pretreatment groups.

## Discussion

Our results demonstrate that distinct intestinal ecosystems are differentially impacted by *C. difficile* colonization and that these changes to community metabolism could have implications for the ability of the pathogen to persist in those environments. Furthermore, our mutli-omics approach demonstrated that *C. difficile* manipulated the niche landscape of the intestinal tract. Instances of active nutrient niche restructuring in the gut have been documented previously for prominent symbiotic bacterial species in gnotobiotic mice (25), but not in a conventionally-reared animal model of infection following antibiotic pretreatment. Interestingly, the taxonomic groups that produced the transcripts that were most altered by *C. difficile* colonization were rare in their cecal community. Previous studies have found that rare taxonomic groups, even those at a low abundance as a result of a spontaneous perturbation, may have disproportionate effects on the metabolome of the rest of the community (26). For example, in temperate lakes conditionally rare microbes were found to be far more metabolically active than highly abundant taxa (27). These examples of response to perturbations are interesting models for thinking about the dynamics of bacterial populations recovering from an antibiotic perturbation. As such, *C. difficile* may compete with these organisms to ultimately affect greater change to the entire ecosystem and open a long-lasting nutrient niche. While this hypothesis requires further exploration, it provides an ecological framework to study the interactions between *C. difficile* and members of susceptible communities.

This study is one of the first *in vivo* observations that a medically relevant bacterial pathogen may alter the metabolic activity of a host-associated community to promote its own colonization. This is also the first application of metatranscriptomic analysis of the gut microbiota *in vivo* and in response to a pathogen. Other groups have identified potential metabolite markers of *C. difficile* infection in patient feces (28), but they were not able to identify associations with changes in community metabolism that were afforded to us by our paired metabolomic and metatranscriptomic analyses. In a recent study, a tick-vectored bacterial pathogen altered the ability of the resident microbiota of the tick by interrupting proper biofilm formation and allowing lasting colonization (29). It was also recently found that bacterial metabolic generalists may be more likely to actively antagonize the growth of other species in an environment that they are colonizing (30). We previously showed that *C. difficile* has a wide nutrient niche-space *in vivo* and most likely utilizes its role as a metabolic generalist to colonize diverse gut microbiomes (19). The ability to simultaneously antagonize the metabolism of surrounding populations in cecal environments that support persistence would explain the more significant shifts in those metatranscriptome. While we acknowledge that this study may not elucidate the specific mechanism by which this interaction occurs, the combined systems analysis strengthens each individual level of observation. When the results from these approaches are combined reveals a clearer understanding of *C. difficile*-related microbial ecology. This research lays the groundwork for a more rationale consideration of the metabolic functionalities of bacterial taxa to consider when attempting to rebuild *C. difficile* colonization resistance across differentially perturbed gut environments.

In spite of consistent results across the different methods we used in this study, several limitations should be noted. First, as with all transcriptomic studies, the relative level of mRNA detected for a given gene does not necessarily reflect the amount of functional protein made by a cell or the post-translational modifications that are required to activate the enzymes. Additionally, due to the low relative abundance of *C. difficile* in these communities, it was necessary for us to pool samples to generate a large number of reads from each group rather than sampling multiple replicates within each group. Greater transcript read abundance per gene allowed for improved survey for the activity of lowly abundant species as well as greater confidence in genes found to be highly transcribed. Although the lack of animal-based replication for the metatranscriptomic data does potentially limit the ability to generalize our results, this approach has been successfully utilized by numerous groups in the past to accurately characterize transcriptionally activity across communities of bacteria (19, 31–33). Furthermore, the metatranscriptomic data were supported by the 16S rRNA gene sequence and metabolomic data which were collected from individual animals. With respect to the metabolomic data, alternative interpretations of the data also exist. For example, we assumed that metabolites, which did not change in concentration between uninfected and infected conditions were not impacted by *C. difficile* colonization. However, it is possible that the metabolism of *C. difficile* itself simply substituted for a function that was already present in the uninfected community. The insights gathered from the metatranscriptomic data suggests that this was unlikely. By leveraging multiple methods to test our hypotheses we were able to mediate the weaknesses of any individual method and present a more unified description of the system than any of the methods on their own.

Our study supports the hypothesis that the gut microbiota of healthy individuals maintains colonization resistance to *C. difficile* by outcompeting the pathogen for preferred nutrient niche space. Ultimately, our results suggest that each susceptible and subsequently infected microbiome may be unique and require specific microbes or functionalities to restore colonization resistance against *C. difficile* in that specific context. Conversely, colonization resistance against *C. difficile* may be the result of contributions by distinct sub-communities of bacteria across each unique resistant gut community. Several studies have attempted to identify single bacterial species or consortia that are able to achieve colonization resistance; however, these efforts have only resulted in partially resistance (34–37). Considering the structure and function of the microbiome is intimately connected to colonization resistance against the *C. difficile*, it has become imperative to understand the ecological factors that allow some gut environments to be persistently colonized while others are not. This research lays the groundwork for future studies to assess context dependent restoration of *C. difficile* colonization resistance and what factors are able to interfere with the ability of *C. difficile* to modify gut ecology to promote clearance.

## Materials and Methods

### Animal care and antibiotic administration

Briefly, approximately equal numbers of male and female conventionally-reared six-to-eight week-old C57BL/6 mice were randomly assigned to each experimental group (genders were housed separately). Nine mice were used in each experimental and control group. They were administered one of three antibiotics; cefoperazone, streptomycin, or clindamycin before oral *C. difficile* infection (Table S1). A detailed description of these animal models was outlined previously (19). A similar experimental design was implemented for gnotobiotic mice and was performed with the University of Michigan Germfree Mouse Center as described previously (19). All animal protocols were approved by the University Committee on Use and Care of Animals at the University of Michigan and carried out in accordance with the approved guidelines from the Office of Laboratory Animal Welfare (OLAW), United States Department of Agriculture (USDA) registration, and the Association for Assessment and Accreditation of Laboratory Animal Care (AAALAC). The protocol license Institutional Animal Care and Use Committee (IACUC) number for all described experiments is PRO00006983.

### *C. difficile* infection and necropsy

On the day of challenge, 1×10^3^ *C. difficile* spores were administered to mice via oral gavage in phosphate-buffered saline (PBS) vehicle. Mock-infected animals were given an oral gavage of 100 ul PBS at the same time as those mice administered *C. difficile* spores. 18 hours following infection, mice were euthanized by CO_2_ asphyxiation and necropsied to obtain the cecal contents. Aliquots were immediately flash frozen for later DNA extraction and toxin titer analysis. A third aliquot was transferred to an anaerobic chamber for quantification of *C. difficile* abundance. The remaining content in the ceca was mixed in a stainless steel mortar housed in a dry ice and ethanol bath. Cecal contents from all mice within each pretreatment group were pooled into the mortar prior to grinding to a fine powder. The ground content was then stored at -80°C for subsequent RNA extraction. For 10-day colonization studies, fresh stool was collected from infected mice each day beginning on the day of infection. Mice were monitored for overt signs of disease and were euthanized after the final stool collection.

### *C. difficile* cultivation and quantification

Cecal samples were weighed and serially diluted under anaerobic conditions with anaerobic PBS. Differential plating was performed to quantify *C. difficile* vegetative cells by plating diluted samples on CCFAE plates (fructose agar plus cycloserine, cefoxitin, and erythromycin) at 37°C for 24 hours under anaerobic conditions (38). Quantification of total *C. difficile* CFU for the 10- day colonization experiments was performed from stool using TCCFAE to measure total *C. difficile* load in these animals over time.

### DNA/RNA extraction and sequencing library preparation

DNA for shotgun metagenomic and 16S rRNA gene sequencing was extracted from approximately 50 mg of cecal content from each mouse using the PowerSoil-htp 96 Well Soil DNA isolation kit (MO BIO Laboratories) and an epMotion 5075 automated pipetting system (Eppendorf). The V4 region of the bacterial 16S rRNA gene was amplified using custom barcoded primers (39). Equal molar ratios of raw isolated DNA within each treatment group were then pooled and ~2.5 ng of material was used to generate shotgun libraries with a modified 10-cycle Nextera XT genomic library construction protocol (Illumina). This was done to mimic the pooling strategy necessary for metatranscriptomic library preparation. Final libraries were pooled at equal molar ratios and stored at -20°C. For RNA extraction, a more detailed description of the procedure can be found in (19). Briefly, immediately before RNA extraction, 3 ml of lysis buffer (2% SDS, 16 mM EDTA and 200 mM NaCl) contained in a 50 ml polypropylene conical tube was heated for 5 minutes in a boiling water bath (40). The hot lysis buffer was added to the frozen and ground cecal content. The mixture was boiled with periodic vortexing for another 5 minutes. After boiling, an equal volume of 37°C acid phenol/chloroform was added to the cecal content lysate and incubated at 37°C for 10 minutes with periodic vortexing. The mixture was the centrifuged at 2,500 x g at 4°C for 15 minutes. The aqueous phase was then transferred to a sterile tube and an equal volume of acid phenol/chloroform was added. This mixture was vortexed and centrifuged at 2,500 x g at 4°C for 5 minutes. The process was repeated until aqueous phase was clear. The last extraction was performed with chloroform/isoamyl alcohol to remove acid phenol. An equal volume of isopropanol was added and the extracted nucleic acid was incubated overnight at -20°C. The following day the sample was centrifuged at 12000 x g at 4°C for 45 minutes. The pellet was washed with 0°C 100% ethanol and resuspended in 200 ul of RNase-free water. Following the manufacturer’s protocol, samples were then treated with 2 ul of Turbo DNase for 30 minutes at 37°C. RNA samples were retrieved using the Zymo Quick-RNA MiniPrep according the manufacturer’s protocol. The Ribo- Zero Gold rRNA Removal Kit (Epidemiology) was then used to deplete prokaryotic and eukaryotic rRNA from the samples according the manufacturer’s protocol (Illumina). Stranded RNA-Seq libraries were made constructed with the TruSeq Total RNA Library Preparation Kit v2, both using the manufacturer’s protocol. Completed libraries were stored at -20°C until time of sequencing.

### High-throughput sequencing and raw read curation

Sequencing of 16S rRNA gene amplicon libraries was performed using an Illumina MiSeq sequencer as described previously (39). The 16S rRNA gene sequences were curated using the mothur software package (v1.36) and OTU-based analysis was performed as described in (19). Genus-level classification-based analysis of 16S rRNA gene sequence data was accomplished using the phylotype workflow in mothur and the full SILVA bacterial taxonomy (release 132). Shotgun metagenomic sequencing was performed in 2 phases. Libraries from mock-infected communities, which were also to be utilized for *de novo* contig assembly, were sequenced using an Illumina HiSeq 2500 on 2×250 paired-end settings and was repeated across 2 lanes to normalize for inter-run variation. *C. difficile-infected* metagenomic libraries were sequenced with an Illumina NextSeq 300 with 2×150 settings across 2 runs to also normalize for inter-run variation. These efforts resulted in an average of 280,000,000 paired raw reads per sample. Metatranscriptomic sequencing was performed on an Illumina HiSeq 2500 with 2×50 settings and was repeated across 4 lanes for normalization and to normalize for technical variation between lans and to obtain necessary coverage (32). This gave an average of 380 million raw cDNA reads per library. Both metagenomic and metatranscriptomic sequencing was performed at the University of Michigan Sequencing Core. Raw sequence read curation for both metagenomic and metatranscriptomic datasets was performed in a two step process. Residual 5’ and 3’ Illumina adapter sequences were trimmed using CutAdapt (41) on a per library basis. Reads were quality trimmed using Sickle (42) with a quality cutoff of Q30. This resulted in approximately 270 million reads per library (both paired and orphaned) for both metagenomic and metatranscriptomic sequencing. Actual read abundances for individual metagenomic and metatranscriptomic sequencing efforts can be found in Table S4.

### Metagenomic contig assembly and gene annotation

Metagenomic contigs were assembled using Megahit (43) with the following settings: minimum kmer size of 87, maximum kmer size of 127, and a kmer step size of 10. Prodigal was utilized to to identify putative gene sequences, and were screened for a minimum length of 250 nucleotides. These sequences were translated to amino acids and the predicted peptides were annotated based on the KEGG protein database (44) using Diamond implementation of BLASTp (45). Peptide-level gene annotations were assigned to the corresponding nucleotide sequence, and genes failing to find a match in KEGG were preserved as unannotated genes. Final nucleotide FASTA files with KEGG annotations were then utilized in the construction of Bowtie2 mapping databases from downstream analyses (46).

### DNA/cDNA read mapping and normalization

Mapping of DNA and cDNA reads to the assemblies was accomplished using Bowtie2 and the default stringent settings (46). Optical and PCR duplicates were then removed using Picard MarkDuplicates (http://broadinstitute.github.io/picard/). The remaining mappings were converted to idxstats format using Samtools (47) and the read counts per gene were tabulated. Discordant pair mappings were discarded and counts were then normalized to read length and gene length to give a per base report of gene coverage. Transcript abundance was then normalized to gene abundance to yield overall level of transcription for each gene. Reads contributed by *C. difficile* were removed from analysis using Bowtie2 against the *C. difficile* str. 630 genome with settings allowing for up to 2 mismatches.X

### Quantification of *in vivo* metabolite relative concentrations

Metabolomic analysis was performed by Metabolon (Durham, NC), for a detailed description of the procedure refer to (19). Briefly, all methods utilized a Waters ACQUITY ultra-performance liquid chromatography (UPLC) and a Thermo Scientific Q-Exactive high resolution/accurate mass spectrometer interfaced with a heated electrospray ionization (HESI-II) source and Orbitrap mass analyzer at 35,000 mass resolution. Samples were dried then reconstituted in solvents compatible to each of the four methods. The first, in acidic positive conditions using a C18 column (Waters UPLC BEH C18–2.1×100 mm, 1.7 um) using water and methanol, containing 0.05% perfluoropentanoic acid (PFPA) and 0.1% formic acid (FA). The second method was identical to the first but was chromatographically optimized for more hydrophobic compounds. The third approach utilized a basic negative ion optimized conditions using a separate dedicated C18 column. Basic extracts were gradient eluted from the column using methanol and water, however with 6.5mM Ammonium Bicarbonate at pH 8. Samples were then analyzed via negative ionization following elution from a hydrophilic interaction chromatography column (Waters UPLC BEH Amide 2.1×150 mm, 1.7 um) using a gradient consisting of water and acetonitrile with 10 mM Ammonium Formate, pH 10.8. The MS analysis alternated between MS and data-dependent MS n scans using dynamic exclusion. The scan range varied slighted between methods but covered 70–1000 m/z. Library matches for each compound were checked for each sample and corrected if necessary.

### Statistical methods

All statistical analyses were performed using R (v.3.2.0) and the vegan package (48). Significant differences of inverse Simpson diversity, CFU, toxin titer, and metabolite concentrations were determined by Wilcoxon signed-rank test with Benjamini-Hochberg correction using a study-wide Type I error rate of 0.05. Undetectable points used half the limit of detection for CFU and toxin statistical calculations. Dynamic time warping was performed with the dtw package in R (49). Random forest was performed using the AUCRF implementation (22) as well as the standard package (50) in R. Distances of outlier points from center line during metatranscriptomic comparisons was accomplished using 2-dimensional linear geometry.

### Data Availability

Pooled and quality trimmed *C. difficile-infected* metatranscriptomes (SRA; PRJNA354635) and 16S rRNA gene amplicon read data (SRA; PRJNA383577) from infection experiments are available through the NCBI Sequence Read Archive. Metagenomic reads and mock-infected metatranscriptomic reads can be found also on the SRA (PRJNA415307). Data processing steps beginning with raw sequence data to the final manuscript are hosted at https://github.com/SchlossLab/Jenior_Metatranscriptomics_mSphere_2018.X

## Acknowledgments

The authors would like to acknowledge Charles Koumpouras for assistance with DNA extractions and metabolomic sample preparation. We would also like to acknowledge members of the University of Michigan Germfree Mouse Center, University of Michigan Sequencing Core, and Metabolon for their assistance in experimental design, execution, and data collection.

## Author Contributions

M.L.J. conceived, designed and performed experiments, analyzed data, and drafted the manuscript. J.L.L. performed experiments, analyzed data, and contributed to the manuscript. V.B.Y. contributed to the manuscript. P.D.S. interpreted data and contributed the manuscript. The authors declare no conflicts of interest.

---

**Supplementary Figure 1.** Conserved markers of *C. difficile* colonization susceptibility in mouse cecal microbiomes. **(A)** NMDS ordination of Bray-Curtis distances of OTU relative abundances between mouse cecal communities that are susceptible to colonization by *C. difficile* and those that are resistant. **(B)** Relative abundance of OTUs included in the optimal model generated by AUCRF classifying the same groups as in panel A, labeled with the finest resolution provided by classifying to the RDP reference database. **(C)** NMDS ordination of Bray-Curtis distances between metabolite intensities. **(D)** Scaled metabolites relative abundnaces included the optimal model generated by AUCRF classifying resistant and susceptible cecal microbiomes. Significant differences in A & C were also calculated using permANOVA. The AUCRF models generated in this analysis also had 0% out of bag error and significant differences in B & D were calculated as in Figure 2.

**Supplementary Figure 2.** Signatures of infection effect on the cecal microbiomes conserved across pretreatment groups. **(A)** NMDS ordination of Bray-Curtis distances of OTU relative abundances between antibiotic-pretreated mouse cecal communities that are either *C. difficile-colonized* or mock-infected. **(B)** Relative abundance of OTUs included the optimal model generated by AUCRF classifying the same groups as in panel A. **(C)** NMDS ordination of Bray-Curtis distances using metabolite intensities between the same classes. **(D)** Scaled intensity of metabolites included the optimal model classifying infected and uninfected cecal microbiomes. Statistical differences were performed as in Figure 2.

**Supplementary Figure 3.** Relative concentrations of select *C. difficile* Stickland fermentation metabolites across infection models. Metabolites included in this analysis were chosen based on their previously published interaction with *C. difficile* Stickland fermentation: **(A)** Proline, **(B)** 4-Hydroxyproline, **(C)** Glycine, **(D)** 5-Aminovalerate. Significant differences were determined by Wilcoxon rank-sum test with Benjamini-Hochberg correction when necessary. Black asterisks in the plotting area represent within group differences, while green asterisks along the top border denote significant differences compared to untreated control.

**Supplementary Figure 4.** Unique genes with functional annotation detectable within each metagenome and metatranscriptome. Genes in each datasets were derived from respective metagenomic assemblies, with only those genes that mapped to a KEGG pathway-level annotation: **(A)** Untreated, **(B)** Streptomycin-pretreated, **(C)** Cefoperazone-pretreated, and **(D)** Clindamycin-pretreated mice. Each panel includes that treatments’ unique genes from metagenomic assembly and genes that recruited at least one cDNA read from the corresponding metatranscriptomes. Collector’s curves from rarefaction analysis of reads mapped to genes from **(E)** metagenomes and **(F)** metatranscriptomes.

**Supplementary Table 1.** Antibiotic pretreatment regime summaries. Antibiotic classes, mechanisms, and dosage information for each pretreatment. Quantified effect on alpha- and beta-diversities of the cecal microbiota are also included.

**Supplementary Table 2.** Summary of impact of infection on cecal community structure and metabolome. Global effect as well as changes to specific metabolites are included.

**Supplementary Table 3.** Summary statistics for datasets containing replicates generated during this study.

**Supplementary Table 4.** High-throughput sequencing read counts and metagenomic assembly. Raw and curated read abundances for both metagenomic and metatranscriptomic sequencing efforts. Raw read curation steps are outlined in Materials & Methods. Metagenomic contig summary statistics reflect the quality of assembly for each group.

